# A method for recognizing motor imagery EEG signals based on high-quality lead selection

**DOI:** 10.1101/2024.03.28.587312

**Authors:** Xiuli Du, Hanxing Wang, Meiya Kong, Meiling Xi, Yana Lv

## Abstract

In the application of motor imagery brain-computer interface system, high-density leads bring redundant noise, which leads to time-consuming system operation and poor performance. A channel selection strategy based on brain function network is proposed. This method introduces Synchronization likelihood was used as a connection index to construct a motor imagery brain functional network, and the centrality analysis of the constructed network was used to select the combination of strong motor-related leads. Experiments were carried out on the EEG datasets dataset IVa of the 3rd International Brain-Computer Interface Competition and dataset I of the 4th International Brain-Computer Interface Competition, and 27 high-quality channels were selected from the 118 channels of dataset IVa, as well as 16 high-quality channels were selected from the 59 channels of dataset I. Finally, the CSP algorithm and support vector machine are used to extract features and classify. The experimental results show that the proposed channel selection strategy can greatly reduce the number of channels while obtaining higher recognition accuracy, which verifies the practicability and effectiveness of the proposed method.

## Introduction

Brain-computer interfaces (BCIs) allow individuals to interact with the real world solely through brain neural activity [1]. Motor imagery electroencephalography (MI-EEG) is one of the widely used paradigms in BCI, primarily applied in the field of motor rehabilitation. To obtain comprehensive EEG information, most EEG acquisition devices utilize high-density electrode arrays to record EEG data [2]. However, multi-channel recording, while offering more EEG information, also introduces noise and additional redundant information, increasing data dimensionality and impacting model performance. Therefore, the selection of effective electrode combinations is crucial for improving classification accuracy [3]. Currently, there are four main algorithms used for electrode selection in EEG signals, namely Filter methods, Wrapper methods, Embedded methods, and Hybrid selection methods.

“Filter methods” use evaluation metrics such as information measures, dependency measures, distance measures, and consistency measures to determine candidate subsets, which are generated using search algorithms [4][5]. In “Filter methods”, electrode selection is independent of the classifier, offering faster computation speed and better flexibility. However, the quality of selected electrodes is often low, resulting in poor EEG signal recognition. “Wrapper methods” evaluate electrode feature subsets based on the classification accuracy of the classifier, continually refining the feature subset based on classification results. While effective, “Wrapper methods” incur high computational complexity and overhead [6]. “Embedded selection methods” refer to electrode selection strategies deeply integrated with the classifier, selecting electrodes during the classifier training process. This approach effectively avoids overfitting during feature selection and incurs low computational complexity [7]. However, it may restrict the choice of classifiers. “Hybrid selection methods” combine “Filter and Wrapper methods”, eliminating pre-specified norms and stopping criteria. Hybrid techniques typically utilize an independent metric and a mining algorithm to evaluate available channel subsets, suitable for larger datasets [8]. Existing algorithms often rely on a single evaluation metric or depend on recognition accuracy for electrode selection. However, the ultimate goal of electrode selection is to serve the identification phase in BCI applications, requiring both electrode selection quality and avoidance of overly complex selection algorithms. For experimental data under the same paradigm, a simple and independent method for selecting an effective set of electrodes for subsequent recognition tasks is preferred in practical BCI applications. Therefore, this paper proposes an electrode selection method based on brain functional networks. After constructing the brain functional network, the topological properties of the network are analyzed to select appropriate topological properties as electrode selection indicators, thereby obtaining effective electrode combinations, reducing data dimensionality and computational complexity, and improving classification accuracy.

## 1 Research Methodology

### 1.1 Building MI Functional Networks Based on Synchronization Likelihood

Brain functional networks represent the mathematical representation of the brain as a complex system, defined by a set of nodes and connections (edges) between nodes. Nodes in the brain functional network typically represent brain regions, while connections represent functional connections, corresponding to the magnitude of temporal correlation in activity and may occur between anatomically non-connected regions. According to measurement standards, functional connectivity may reflect linear or nonlinear interactions, as well as interactions at different time scales[9]. The construction process of brain functional networks may include the following steps: selecting reasonable brain regions or signal recording locations as nodes; defining the correlations between nodes as edges; and selecting appropriate thresholds to generate the connectivity network.

#### 1.1.1 Node Definition

Traditionally, the most intuitive nodes would be neurons of the nervous system, considering each neuron as a node and defining synapses connecting neurons as edges. However, this node and edge definition is only suitable for small-scale biological systems of simple organisms within the nervous system. For instance, Bassett [10] constructed a network of the nematode nervous system, considering neurons as nodes, forming a network with 300 nodes and 7600 edges. However, such an approach has clear limitations for complex biological nervous systems like humans. The human cerebral cortex contains over 22 billion neurons, each connected to approximately more than 10,000 other neurons via synapses [11], making it exceedingly challenging for current computational systems. Therefore, in current studies of human brain connectivity, research based on neurons or clusters of neurons as units for network construction remains unattainable, and existing studies are based on large-scale (brain region) node partitions.

The use of EEG sensors provides high-resolution time series, offering a reliable node selection scheme for network construction. Based on EEG collected using the 10-20 EEG recording system, which reflects electrophysiological signals of neural activity within the brain externally, the placement rules are defined by the International Federation of Clinical Neurophysiology, conforming to node definition standards. Thus, when constructing EEG brain networks, scalp electrodes are commonly used as nodes. The scale of nodes varies according to the standards of the international 10-20 EEG recording system, typically including different numbers of channels such as 32, 64, 128, and 256 [12]. Therefore, in this study, the electrode placement positions specified in the dataset being used are employed as nodes for research purposes.

#### 1.1.2 Synchronization Likelihood

In EEG signal research, synchronization phenomena are key features of information exchange between different brain regions. Therefore, synchrony becomes the primary measure for studying correlations between EEG channels [13]. Commonly used measures of synchrony include cross-correlation, coherence, mutual information (MI), phase synchronization (PS), synchronization likelihood (SL), and phase lag index (PLI). Metrics such as coherence, phase synchronization, and phase lag index describe linear correlations. However, EEG is a stochastic nonlinear time series, containing numerous nonlinear features. Using linear correlation metrics cannot fully characterize its nonlinear synchronization relationships. Synchronization likelihood is a novel method for exploring nonlinear dependencies and generalized synchrony in multivariate datasets, possessing good representation capabilities for both linear and nonlinear features in EEG signals, and exhibiting robustness to nonlinear time series [14]. Therefore, this study adopts SL as the metric for constructing motor imagery functional brain networks.

Synchronization likelihood reconstructs each time series into a series of specific “patterns” and then detects the regularity of simultaneous occurrence of these “patterns.” The algorithm for synchronization likelihood is described as follows:

1. For each electrode time series *X*_*k,i*_ perform phase space reconstruction:

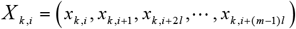

In the equation, *k* represents the number of channels in the collected signal, *I* denotes discrete time points, *l* is the delay time, and _*m*_ is the embedding dimension.
2. For the time series of channel *k* and time points *i*, the probability that the distance between embedding vectors is less than distance_*ε*_ is defined as:

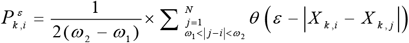

In the equation, |·| represents the Euclidean distance, and *θ* is the Heaviside step function. When *x* ≤0 时, *θ*(*x*) = 0; When *x* > 0,*θ*(*x*) =1 ; *ω*_1_, *ω*_2_ denote two time windows, where *ω*_1_ is the Theiler correction for autocorrelation effects, *ω*_2_ sharpens the time resolution of synchronization measurement, and they satisfy *ω*_1_ ≤*ω*_2_ ≤ *N*.
3. For each channel *k* and time point *i*, determine its critical distance*ε*_*k,i*_, such that 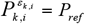, where, *P*_*ref*_ = 1.
4. Determine the number of channels for each discrete time pair (*i, j*), where the distance between embedding vectors *X*_*k,i*_ and *X* _*k, j*_ in the window *ω*_1_ < |*j* − *i*| < *ω*_2_ is less than the critical distance:

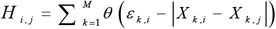

Where, *H*_*i, j*_ ranges from 0 to *M*.
5. For each channel *k* and discrete time pair (*i, j*), define the synchronization likelihood *S*_*k,i, j*_ as:

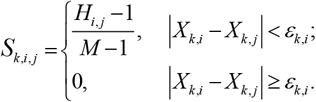

Take the average over all *j* to obtain the synchronization likelihood *S*_*k, i*_

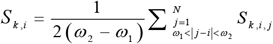

In the equation, *S*_*k, I*_ measures the strength of synchronization of channel *k*at time point *i* with the other *M*−1 channels. The value of *S*_*k, i*_ ranges between *P*_*ref*_ and 1 之间. *S*_*k, i*_ = *P*_*ref*_ indicates that all *M* time series are uncorrelated, while *S*_*k,i*_ =1 indicates maximum synchronization among all *M* time series.

#### 1.1.3 Constructing the MI Brain FunctionalNetwork

The main steps for constructing the MI functional brain network are as follows:

1. Define the nodes of the network: For multi-channel EEG, typically consider *k*EEG electrodes as nodes.
2. Define the edges of the network: Select synchronization likelihood to measure the functional connections between nodes, thus obtaining an association matrix of the functional connection strength between all pairs of nodes.
3. Select appropriate thresholds: Threshold the connectivity strength matrix to remove spurious connections, resulting in a binary matrix *Q*.

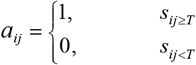

In this context, where *a*_*ij*_ = 1 indicates the presence of a connection edge between nodes *i* and *j*, and *a*_*ij*_ = 0 indicates the absence of a connection edge, the topological structure of the MI functional brain network can be derived from the matrix *Q*.

### 1.2 Based on Brain Functional Network, Selecting Electrode Combinations

#### 1.2.1 Network Centrality Metrics

Important nodes in a network refer to a few nodes that, compared to other nodes in the network, can significantly influence the network’s topological structure and functional connections. Although the number of important nodes in a network is small, their influence can rapidly propagate to most nodes in the network. Various centrality metrics have been developed to identify important nodes in networks. Degree centrality and betweenness centrality, as the most fundamental and important metrics, are widely used in the selection of important nodes.

Node degree refers to the total number of edges connected to that node. The higher the node degree, the more important the node’s position in the network. The average node degree in the network is the average value of the node degrees calculated over *k* nodes. By normalizing the node degree, we obtain degree centrality[16]. Degree centrality measures the importance of individual nodes as intersections of multiple connections in a functional network and is the most direct indicator of centrality. The definitions of node degree, average node degree, and degree centrality are as follows:

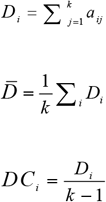

In the equation, *a* _*i j*_ represents the elements of the binary network, *k* denotes the number of nodes, *D*_*i*_ represents the degree of a node, *DC*_*i*_ indicates the degree centrality of a node, and 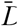 represents the average node degree of the network.

Betweenness centrality is another indicator characterizing the centrality of nodes. It represents the proportion of the shortest paths in the functional network that pass through a particular node [17]. The higher the betweenness centrality of a node, the more information flow it carries, indicating a heavier load, and consequently, a greater impact on the performance of the motor imagery brain network. Betweenness centrality is defined as:

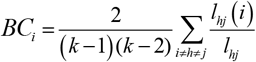

In the equation, *k* represents the number of nodes, *l*_*hj*_ denotes the characteristic path length, and *l*_*hj*_ (*i*) represents the characteristic path length passing through node *i*.

#### 1.2.2 Selection of Electrode Combinations

Degree centrality reflects the connectivity of each node with other nodes, representing local characteristics. Betweenness centrality, on the other hand, characterizes the importance of nodes based on the number of shortest paths passing through a particular node, reflecting global characteristics. The influence of these two measures on network-important nodes comes from different aspects. By setting the weight factors of both measures to 50%, we compute a filtering factor *F*_*i*_, the magnitude of *F*_*I*_ reflects the importance of node *i* 节 in the network.

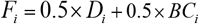

*F*_*i*_ values are sorted in descending order, and nodes with lower *F*_*i*_ values are removed, while the remaining nodes are retained. This process is repeated for all participants, and the remaining nodes across all participants are compared. The nodes covered by all participants are selected as the final filtering result, thereby completing the selection of highly correlated electrode combinations for motor imagery. Considering the current results of electrode selection methods, it is evident that preserving one-fourth of the total number of electrodes can effectively retain EEG information [5] [6][18][19]. Therefore, in this study, when removing nodes with lower *F*_*i*_ values, a strategy of removing half of the nodes is adopted. Specifically, the bottom 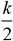 nodes based on the *F*_*I*_ ranking are discarded, and the remaining half is retained based on the coverage by participants, resulting in the optimal electrode combination.

## 2 Experimental Results Analysis

### 2.1 Experimental Data

#### Dataset One

The experiment utilized data from the Fourth International Brain-Computer Interface Competition held in 2008 (BCI competition IV dataset I). The dataset consists of EEG data from seven participants, recorded using 64 electrodes (with 59 effective channels), distributed according to the international 10-20 system. Since the data from participants c, d, and e were artificially generated, this study focused on EEG data from participants a, b, f, and g. The data include tasks involving motor imagery of the left hand, right hand, and both feet, with each trial involving only two types of motor imagery. The experimental paradigm for a single motor imagery task is depicted in Figure 1: participants sit in front of a screen, with the computer screen blank at the start of the trial, while the participants remain in a relaxed state. At the 2nd second, a brief beep sound is provided to indicate that the participant should prepare to start. From the 2nd to the 6th second, arrows pointing left, right, or down (corresponding to tasks involving imagery of the left hand, right hand, and feet, respectively) appear on the screen, and participants perform the corresponding motor imagery task based on the direction indicated by the arrows. Participants continue to perform the motor imagery task until the crosshair on the screen disappears at t = 6s. After a brief rest with the screen blank again, the next trial commences. The experiment consists of a total of 200 imagery tasks, with a sampling frequency of 100Hz. Only the main rhythmic components associated with movement, namely mu rhythm (8-12Hz) and beta rhythm (15-28Hz), were required. Therefore, the data were band-pass filtered from 8 to 28Hz.

**Figure 1:**
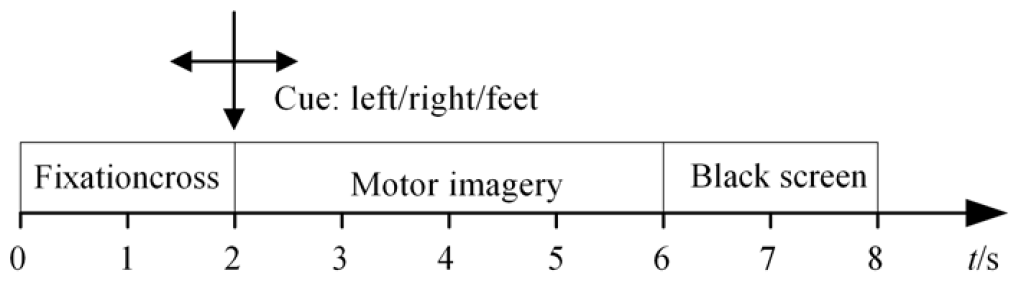
Experimental Paradigm of BCI Competition IV Dataset I

#### Dataset One

The second dataset utilized EEG data from the Third International Brain-Computer Interface Competition held in 2005 (BCI competition III dataset IVa). The dataset consists of EEG data from five healthy participants, recorded using a 118-channel electrode cap with electrodes placed according to the international 10-20 system. The five healthy participants are labeled as aa, al, av, aw, and ay. The data include tasks involving motor imagery of the left hand, right hand, and feet. The experimental paradigm for a single motor imagery task is illustrated in Figure 2: visual cues last for 3.5 seconds, during which participants perform tasks involving imagery of the left hand, right hand, and feet. Subsequently, participants relax for an unspecified duration ranging from 1.75 to 2.25 seconds. The data include time markers for 280 trials for each participant. The original EEG sampling frequency was 1000Hz, which was subsequently downsampled to 100Hz. Similar to the first dataset, only the main rhythmic components associated with movement, namely mu rhythm (8-12Hz) and beta rhythm (15-28Hz), were required. Therefore, the data were band-pass filtered from 8 to 28Hz.

**Figure 2:**
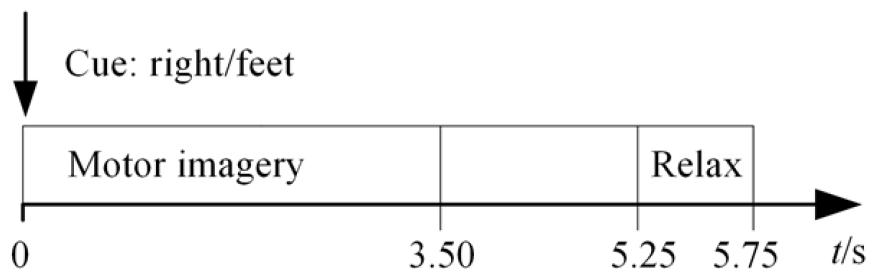
Experimental Paradigm of BCI Competition III Dataset Iva

**Figure 3:**
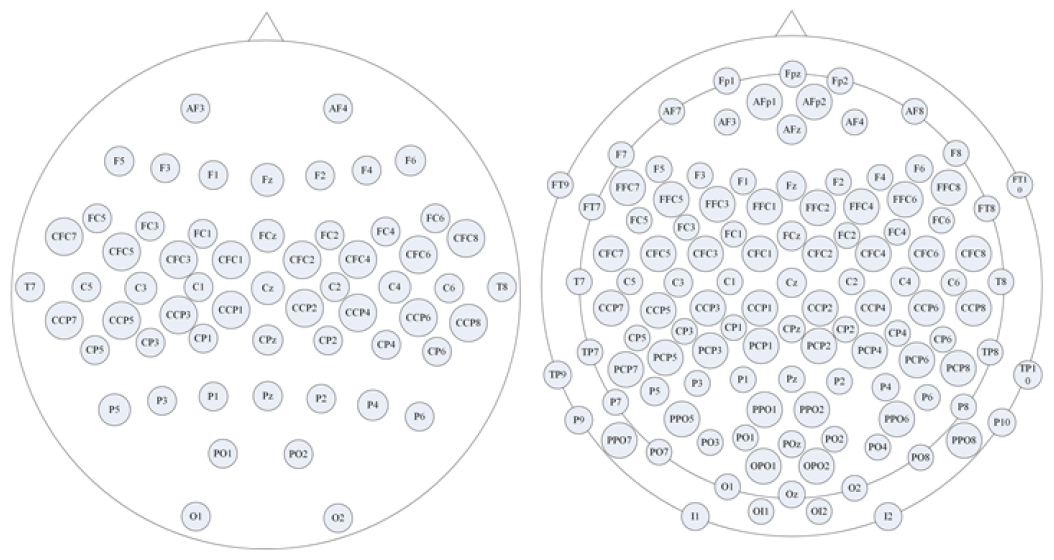
Distribution of Electrode Placements for Dataset I and Dataset IVa

### 2.2 Brain Functional Network Construction and Electrode Selection

Using *k* EEG electrodes as nodes and synchrony likelihood as the metric for functional connectivity between nodes, we obtained an association matrix representing the functional connectivity strength between each pair of nodes. Subsequently, we constructed the MI brain functional network by thresholding the association matrix to remove spurious connections. There is no established standard for selecting the threshold in the process of constructing functional brain networks. The nervous system exhibits characteristics of a small-world network. To ensure that a graph with *k* nodes is fully connected, the average node degree of the network should be greater than or equal to the natural logarithm of the number of network nodes, denoted as 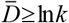 [20]. The ultimate goal of this study is to select electrode combinations based on centrality metrics, and further centrality analysis is needed in the next step. Therefore, the threshold selection principle here is to remove only spurious links and connections with very weak connectivity..

When the threshold is too small, many false connections may exist, resulting in a decrease in the small-world properties of the brain network, making it unable to reflect important information. Conversely, when the threshold is too large, isolated nodes may appear in the network, leading to a decline in the functional connectivity of the network, and the integrity of brain network information cannot be guaranteed. Figure 5 shows the distribution of the average node degree 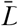 characteristics for both datasets with thresholds increasing in increments of 0.05.

**Figure 5:**
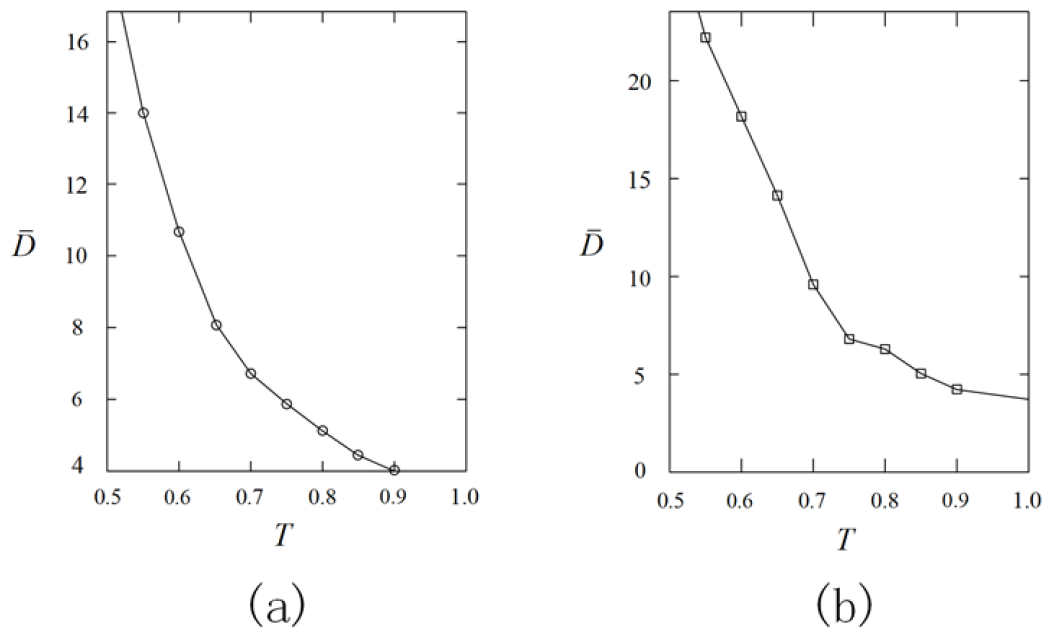
Distribution of Thresholds and Average Node Degrees Characteristics

For dataset one, as shown in Figure 5(a), when the threshold is set to 0.65, the range of the average node degree is 7 to 9, satisfying 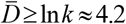;For dataset two, as depicted in Figure 5(b), when the threshold is set to 0.7, the range of the average node degree is 8 to 11, also satisfying 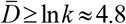. At this point, the MI brain functional networks exhibit higher small-world properties, and the integrity of network information is ensured.

Figure 6 depicts the constructed topology of the brain functional networks, where (a) represents the MI brain functional network topology of participant a in dataset one, and (b) represents the MI brain functional network topology of participant aa in dataset two. From Figure 6, it can be visually observed that the strongly connected electrodes are mainly distributed in the central posterior and central anterior regions of the brain, corresponding to the motor sensory areas, which align with physiological cognition.

**Figure 6:**
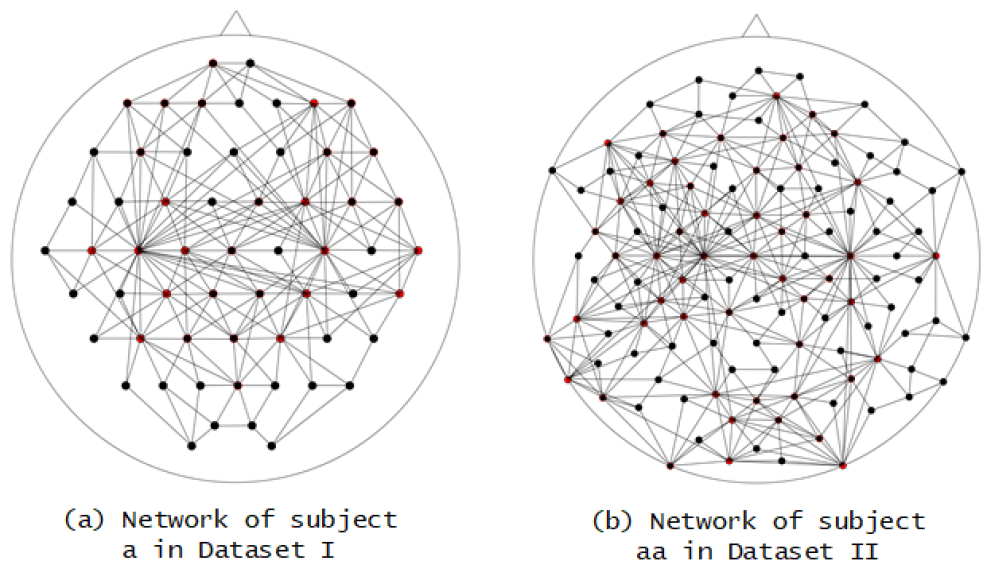
Topological Structure of Brain Functional Networks

The experimental results of this study demonstrate that although each participant exhibits different functional connectivity patterns during the imagination task, the activated electrode nodes ultimately distribute across the motor sensory area. Following the construction of functional brain networks for all participants, the corresponding selection factor *F*_*i*_ was computed based on the node degree centrality and betweenness centrality indices. Then, the lower half of nodes ranked by *F*_*i*_ values were removed. Finally, the retained nodes across all participants were compared, and the final combination was selected based on coverage conditions, thus completing the selection of highly correlated imagination-associated electrode combinations. Table 1 presents the final selection information obtained from experiments conducted on all participants in Dataset I and Dataset II. Through comparison, the selected electrode combinations are found to be distributed in the motor sensory areas of the brain, consistent with neurophysiological knowledge.

**Table 1.** Selection Information of Channels Based on Network Centrality Analysis.

| Dataset | Electrode combinations |
| --- | --- |
| Dataset one<br>( BCIC IV dataset I ) | C3、C1、C5、CFC3、CP3、<br>CCP3、F3、FC1、Cz、CPz、<br>C4、CCP4、CP2、CCP8、<br>CFC4、F4 |
| Dataset two<br>( BCIC III dataset<br>IVa ) | C3、C1、CFC5、FC1、F3、<br>FC5、F7、CP3、CP5、TP7、<br>PCP3、P7、PO1、OPO1、<br>FCz、Cz、CPz、C4、CFC4、<br>FFC6、AFp2、T8、CCP2、<br>CCP6、CP4、PCP4、P6 |

### 2.3 Validation of Electrode Combinations

The classic Common Spatial Patterns (CSP) algorithm simultaneously diagonalizes the covariance matrices of two types of signals, maximizing the differences between them. It is commonly used for feature extraction in EEG signal binary classification tasks. Therefore, in this experiment, the CSP algorithm was utilized for feature extraction, with Support Vector Machine (SVM) employed as the classifier, using 10-fold cross-validation. The recognition accuracies for Dataset I (BCIC IV dataset I) and Dataset II (BCIC III dataset IVa) are shown in Tables 2 and 3, respectively. Here, “AC” denotes the usage of all electrodes for motor imagery recognition tasks, while “3C” indicates the use of three commonly selected motor-related electrodes (i.e., C3, Cz, and C4) for motor imagery recognition tasks.

**Table 2:**
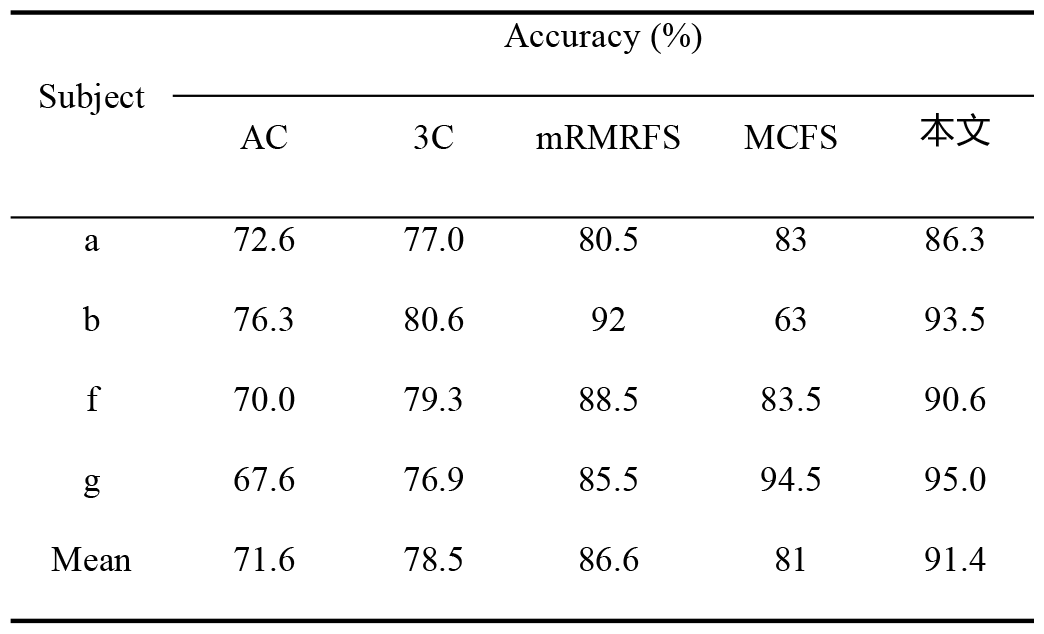
Recognition Accuracy under Different Electrode Selection Strategies for Dataset I.

| Subject | Accuracy (%) |  |  |  |  |
| --- | --- | --- | --- | --- | --- |
|  | AC | 3C | mRMRFs | MCFS | 本文 |
| a | 72.6 | 77.0 | 80.5 | 83 | 86.3 |
| b | 76.3 | 80.6 | 92 | 63 | 93.5 |
| f | 70.0 | 79.3 | 88.5 | 83.5 | 90.6 |
| g | 67.6 | 76.9 | 85.5 | 94.5 | 95.0 |
| Mean | 71.6 | 78.5 | 86.6 | 81 | 91.4 |

**Table 3:** Recognition Accuracy under Different Electrode Selection Strategies for Dataset IVa.

| Subject | Accuracy (%) |
| --- | --- |

|  | AC | 3C | CSP-RMF | MCFS | 本文 |
| --- | --- | --- | --- | --- | --- |
| aa | 65.6 | 66.8 | 81.4 | 80.7 | 84.9 |
| al | 85.0 | 79.3 | 92.4 | 97.1 | 95.8 |
| av | 69.1 | 74.2 | 70.0 | 72.9 | 82.9 |
| aw | 86.3 | 89.5 | 83.6 | 93.2 | 96.4 |
| ay | 74.0 | 73.9 | 85.0 | 92.9 | 95.6 |
| Mean | 76.0 | 76.7 | 82.5 | 87.4 | 91.1 |

In Table 2, mRMRFS refers to an algorithm proposed by Chen Shuli [5] et al., which utilizes mutual information for forward-search electrode selection, achieving an average recognition accuracy of 86.6%. MCFS, proposed by Yin Feiyu [4] et al., is a forward-search algorithm based on multi-correlation, achieving an average recognition accuracy of 81%. The strategy proposed in this paper, which selects electrodes based on brain functional network centrality indices, ultimately achieves a recognition accuracy of 91.4%, surpassing the mRMRFS method by 4.8 percentage points and the MCFS method by 10.4 percentage points. In Table 3, CSP-R-MF refers to a method proposed by Feng [18] et al., which is a multi-frequency band EEG (CSP-RMF) approach based on common spatial pattern (CSP) channel selection, achieving an average recognition accuracy of 82.5%. MCFS, proposed by Yin Feiyu et al., achieves an average recognition accuracy of 87.4% [4]. The method proposed in this paper ultimately achieves a recognition accuracy of 91.1%, surpassing the CSP-RMF method by 8.6 percentage points and the MCFS method by 3.7 percentage points. These experimental results confirm the scientific validity of the electrode selection strategy proposed in this paper, providing technical reference for the application of BCI systems based on motor imagery..

## 3 Conclusion

In BCI applications, selecting channel data relevant to the task is essential to achieve optimal results. Adding irrelevant channels introduces noise interference, slowing down system operation and reducing recognition performance. Inspired by complex network analysis, this study conducts topological property analysis based on brain functional networks, and selects channel combinations strongly correlated with motor imagery tasks. Specifically, synchrony likelihood is introduced as a connectivity metric to construct motor imagery brain functional networks. Subsequently, the degree centrality and betweenness centrality of the constructed networks are analyzed, and filtering factors are obtained based on centrality metrics. Then, non-important nodes are removed according to the filtering factors, and the final motor imagery strongly correlated channel combinations are obtained based on the coverage of important nodes by all participants. Experiments were conducted on EEG datasets dataset IVa of the third International Brain-Computer Interface Competition and dataset I of the fourth competition. From the 118 channels in dataset IVa, 27 high-quality channels were selected, and from the 59 channels in dataset I, 16 high-quality channels were selected. Finally, CSP algorithm and support vector machine were used for feature extraction and classification recognition of the selected channel data. The experimental results show that the proposed channel selection strategy significantly reduces the number of channels while achieving high recognition accuracy, verifying the practicality and effectiveness of the proposed method.

## References

[1] Raj A, Kumar A. Brain Computer Interface[J]. 2015.

[2] Stam C J, Dijk B. Synchronization likelihood: an unbiased measure of generalized synchronization in multivariate data sets[M]. Elsevier Science Publishers B. V. 2002.

[3] Alani A, Alsukker A. Effect of feature and channel selection on EEG classification.[C]//International Conference of the IEEE Engineering in Medicine & Biology Society. Conf Proc IEEE Eng Med Biol Soc, 2006.

[4] YIN Feiyu, JIN Jing, WANG Xingyu. Channel Selection Based on Multi-Correlation Forward Searching Algorithm for MI Classification[J]. Journal of East China University of Science and Technology, 2020, 46(6): 792–799. DOI: 10.14135/j.cnki.1006-3080.20190901002

[5] hen Shuli, Li Xinjian, Hu Yuxia, Lu Peng, Zhang Rui. Forward searching channel selection algorithm based on mutual information for brain-computer interface [J]. Application Research of Computers,2018,35(04):1080–1083+1087.

[6] Pacheco A F, Nekovee M, Moreno Y. Dynamics of Rumor Spreading in Complex Networks:, 10.1103/PhysRevE.69.066130[P]. 2003.

[7] Zhang Deming, Yin Guodong, Jin Xianjian, Zhuang Weichao. Two-stage and bi-direction feature selection method for EEG channel based on CSP and SFFS-SFBS[J]. Journal of Southeast University (Natural Science Edition),2019,49(1):125–132. doi:10.3969/j.issn.1001-0505.2019.01.018

[8] Wang Shengyu. Recognition Method of Multi-Task Motor Imagery for Brain-Computer Interface[D]. Beijing Jiaotong University,2020. DOI:10.26944/d.cnki.gbfju.2020.001351.

[9] Zhou, D., Thompson, W.K., Siegle, G., 2009. Matlab toolbox for functional connectivity. Neuroimage. 2009 Oct 1;47(4):1590–160

[10] Bassett D S, Greenfield D L, Meyer-Lindenberg A, et al. Efficient Physical Embedding of Topologically Complex Information Processing Networks in Brains and Computer Circuits[J]. Plos Computational Biology, 2010, 6(4):e1000748.

[11] Kutter E F, Bostroem J, Elger C E, et al. Single Neurons in the Human Brain Encode Numbers[J]. Neuron, 2018.

[12] Acharya J N, Hani A, Cheek J, et al. Guideline 2: Guidelines for Standard Electrode Position Nomenclature.

[13] Cao Rui. Nonlinear and Complex Network Theory in the Application of EEG Data Analysis Research[D]. Taiyuan University of Technology, 2014.

[14] Stam C J, Dijk B. Synchronization likelihood: an unbiased measure of generalized synchronization in multivariate data sets[M]. Elsevier Science Publishers B. V. 2002.

[15] Achard S, Bullmore E. Efficiency and cost of economical brain functional networks [J]. PLoS computational biology, 2007, 3(2): e17.

[16] Vitali S, Glattfelder J B, Battiston S. The Network of Global Corporate Control[J]. Business & Politics, 2011, 15(3):357–379.

[17] Newman M. A measure of betweenness centrality based on random walks[J]. Social Networks, 2003, 27(1):39–54.

[18] Feng J K, Jin J, Daly I, et al. An Optimized Channel Selection Method Based on Multifrequency CSP-Rank for Motor Imagery-Based BCI System[J]. Computational Intelligence and Neuroscience, 2019, 2019(6):1–10.

[19] Barachant A, Bonnet S. Channel Selection Procedure using Riemannian distance for BCI applications[C]//IEEE. IEEE, 2011.

[20] Fan Lingxiao, Gao Yunyuan, Ma Yuliang. Classification of Motor Imagery EEG Based on Improved Time-Frequency Common Space Pattern[J]. Journal of Transduction Technology,2022,35(11):1483–1490.

[21] Jin J, Miao Y, Daly I, et al. Correlation-based channel selection and regularized feature optimization for MI-based BCI[J]. Neural Networks, 2019, 118

